# 10β-Hydroxyestra-1,4-diene-3,17-dione Does Not Bind to the Nuclear Estrogen Receptor α

**DOI:** 10.1101/2022.08.04.501604

**Authors:** Katalin Prokai-Tatrai, Laszlo Prokai

## Abstract

The lack of nuclear estrogen receptor (ERα and ERβ) bindings of 10β-hydroxyestra-1,4-diene-3,17-dione (HEDD) and structurally related steroidal *para*-quinols have been shown by an extensive series of multidisciplinary investigational evidence including specific receptor binding studies. In support of the latter, the absence of estrogen-derived *para*-quinols’ in vivo uterotrophic effects has also been well documented. Via in silico docking, a recent publication by Canário et al. (2022) reported a robust binding of HEDD (Figure 1B) to ERα. The authors claimed a strong binding of HEDD — as strong as that of its natural ligand, 17β-estradiol (E2), the main human estrogen. However, an examination of the virtual binding pocket revealed that at least one residue near the critical ligand-binding site of their reported HEDD–ERα complex was labelled as “unknown” indicating thereby alteration of the receptor’s published structure (Tannenbaum et al, 1998; Bafna et al., 2020) to fit the ligand. Based on these arguments, the contradictory result by Canário et al. (2022) on HEDD’s binding to ERα should be dismissed.

## INTRODUCTION

We have previously reported our discovery of the unique and distinguishing properties of HEDD (Prokai et al., 2003). These properties include the ability to serve as a first-in-class central nervous system (CNS-)selective bioprecursor prodrug of estrone (E1, Figure 1A). HEDD, originally termed E1-quinol before we introduced its acronym based on its chemical name (Prokai-Tatrai and Prokai, 2019), must be devoid of estrogenic activities to qualify as an inert prodrug of the hormone. Therefore, HEDD should not exhibit appreciable binding to the nuclear estrogen receptors (ERα and ERβ) and consequently should not trigger estrogen-sensitive tissue proliferations associated with ERα activation.

**Figure 1.**
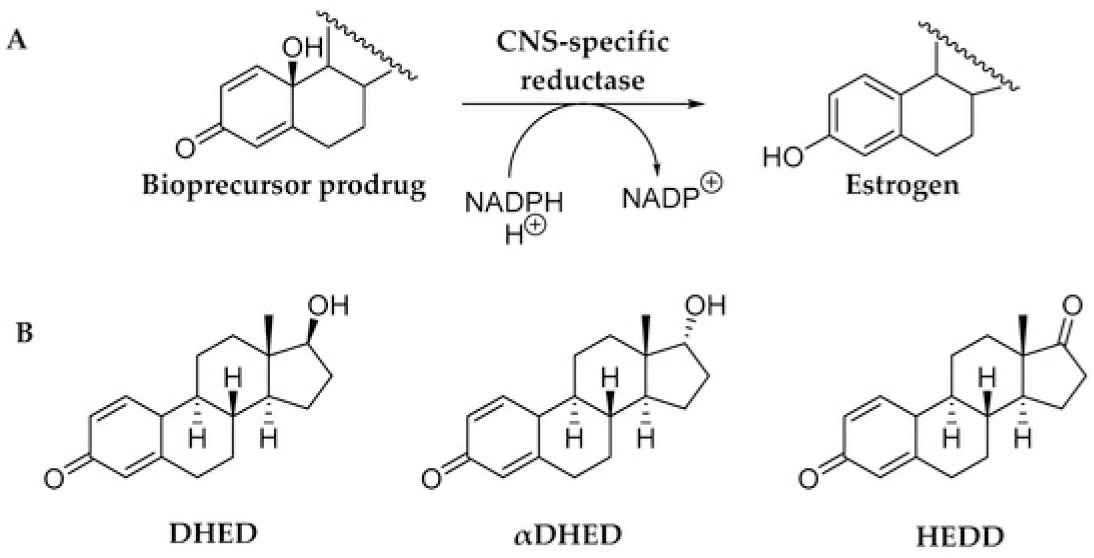
(**A**) Schematic illustration of CNS-selective reductive bioactivation of bioprecursor prodrugs shown in panel B to the corresponding estrogen (E2, αE2 or E1). (B) Chemical structures of bioprecursor prodrugs of estrogens: 10β,17β-dihydroxyestra-1,4-dien-3-one (DHED) for E2; 10β,17α-dihydroxyestra-1,4-dien-3-one (αDHED) for αE2, and 10β-hydroxyestra-1,4-dien-3,17-dione (HEDD) for E1 (Prokai-Tatrai and Prokai, 2018).

## Results and Discussion

Competitive estrogen-receptor (ER) binding studies summarized in Table 1 revealed that HEDD and the structurally closely related E2-derived *para*-quinol 10β,17β-dihydroxyestra-1,4-dien-3-one (DHED), as well as its epimer 10β,17α-dihydroxyestra-1,4-dien-3-one (αDHED) shown in Figure 1B lack appreciable bindings to ERs (Prokai et al., 2003, 2015; Prokai-Tatrai et al., 2018). It is not surprising, because the importance of the phenolic hydroxyl group in ER bindings has been well-established (Anstead et al., 1997; Paterni et al., 2014; McCullough, et al., 2014; Bafna et al., 2020). For example, blocking the phenolic hydroxyl group by ether formation or shielding it by bulky substituents abolishes ER binding and, thus, estrogenicity of the otherwise estrogenic compounds (Prokai-Tatrai et al., 2016).

**Table 1.** Lack of ER-binding affinity of HEDD, DHED and αDHED bioprecursor prodrugs. The ER bindings of their corresponding parent estrogen (E1, E2 and αE2, respectively) are shown as references (Prokai et al., 2003, 2015; Prokai-Tatrai et al., 2018).

| Test Agent | Receptor Binding (IC<sub>50</sub> nM) |  |
| --- | --- | --- |
| | ERα | ERβ |
| E1 | 6.1 | 3.5 |
| HEDD | > 10<sup>4</sup> | > 10<sup>4</sup> |
| E2 | 1.3 | 0.7 |
| DHED | > 10<sup>4</sup> | > 10<sup>4</sup> |
| αE2 | 2.56 | 16.2 |
| αDHED | > 10<sup>4</sup> | > 10<sup>4</sup> |

Through an extensive series of supporting and multidisciplinary experiments as well as validated bioanalytical assays (e.g., Prokai et al, 2003, 2015; Prokai-Tatrai et al., 2018, 2020, 2021; Merchenthaler et al., 2016, 2020), metabolism of these steroidal *para*-quinol bioprecursor prodrugs (HEDD, DHED and αDHED) to the corresponding estrogen (E1, E2 and 17α-estradiol [αE2], respectively) has been unequivocally demonstrated to occur in the CNS (Figure 1A) without peripheral hormonal effects. Specifically, these experiments explicitly revealed that estrogen-sensitive organs such as the uterus, seminal vesicle and anterior pituitary were not stimulated by HEDD, DHED or αDHED treatments, even after chronic dosing. These observations are in complete agreement with the absence of physiologically relevant ER binding of these steroidal *para*-quinols (Table 1), specifically regarding ERα whose role has been well-established in the context of estrogen-induced tissue proliferation (Anstead et al., 1997; Paterni et al., 2014; Winuthayanon et al., 2014).

Accordingly, the lack of uterotrophic effect (i.e., lack of uterus weight gain) by HEDD (Figure 2) or other structurally closely related steroidal *para-*quinols (DHED and αDHED) is a robust surrogate marker for the absence of ERα binding in vivo, which further substantiates receptor binding studies summarized in Table 1.

Contradicting and disregarding these published experimental data yet adopting our HEDD acronym, Canário et al. (2022) has recently asserted via their in silico docking study a strong binding of HEDD to ERα (–10.1 kcal/mol), which would essentially indicate binding as strong as that of ERα’s natural co-crystallized ligand E2, the main human estrogen, with a binding free energy of −9.9 kcal/mol. This would imply that ERα was highly tolerant to the replacement of the phenolic A-ring of E2 with a non-aromatic *para*-quinol in HEDD, disputing the published crystal structure of ERα-E2 complex (Tannenbaum et al., 1998) and extensive pharmacophore studies in this regard.

Nevertheless, our attempts to reproduce HEDD’s docking to ERα using the Autodock Vina protocol and parameters provided by Canário et al. (2022) failed to generate a docked pose with the purported binding energy of the bound complex. A close examination of their virtual binding pocket also revealed that at least one residue was labelled unknown (“UNK1”) near the critical ligand-binding site of their reported HEDD–ERα complex, which indicated an unspecified alteration of the published receptor’s structure. As the latter has been well known (Tannenbaum et al, 1998; Bafna et al., 2020), it should not contain an “unknown” amino acid residue upon proper modelling.

Additionally, the authors’ docking of HEDD to aromatase should also be considered highly irrelevant in a biochemical context, because Canário et al. (2022) should have been aware that conversion of a *para*-quinol to the corresponding estrogen is a reductive process, while aromatase produces estrogens from androgens by a three-step oxidative process (Yoshimoto and Guengerich, 2014; Prokai et al., 2013; Prokai-Tatrai et al., 2018).

## CONCLUSION

Altogether, an extensive series of multidisciplinary and supporting experimental data, including specific receptor binding studies (Table 1), has revealed that HEDD, DHED and αDHED are inert, and indeed are CNS-selective prodrugs of the corresponding estrogens. Therefore, we firmly uphold that these estrogen-derived *para*-quinols are not ligands of ERs. For Canário et al. (2022), the published investigational evidence showing lack of HEDD’s affinity to ERα should have served as a starting point and a quality control for *in silico* docking attempts that without a doubt resulted in their artefactual prediction of binding.

## METHODS

### Synthesis

HEDD and DHED were synthesized by the stereoselective oxidation of the corresponding estrogen (E1 and E2, respectively) in the presence of benzoyl peroxide as a radical initiator and under light irradiation in refluxing dry dichloromethane (Prokai et al., 2003, 2015). Microwave-assisted oxidation of αE2 to produce αDHED was done according to the published protocol (Prokai-Tatrai, 2018).

### ER Binding Studies

Competitive ER-binding binding assays for HEDD and DHED were performed using the HitHunter (Fremont, CA) enzyme-fragment complementation estrogen chemiluminescence assay kit as described before (Prokai et al., 2003). The corresponding estrogens (E1 and E2) served as positive controls. Briefly, 5 nM recombinant ERα or ERβ (Panvera, Madison, WI) and E2-conjugated enzyme donor for 1.5 h were incubated at ligand concentrations ranging from 10 pM to 10 μM. Then, the enzyme acceptor was added to incubate for another 1.5 h, after which chemiluminescence substrate was added for another hour of continued incubation. Relative luminescences were determined by using a Biotek FL600 plate reader (Biotek Instruments, Winooski, VT). A 4-parameter logarithmic curve fitting was used to determine IC_50_ values. Affinities of αE2 and αDHED to ERs were measured using the LantaScreen TR-FRET competitive binding assay screening protocol by the SelectScreen Biochemical Nuclear Receptor Profiling Service (Thermo Fisher Scientific, Waltham, MA).

### Uterotrophic Effect

For the assay, overiectomized (OVX) rats or mice are used. These animals are commercially available from the suppliers or the surgery is done in-house. Approximately three weeks following ovariectomy, animals are randomly assigned to the treatment groups. At predetermined time following the last treatment, animals are euthanized. The uteri are removed from the body cavity, trimmed of excess fat, and connective tissues and the tissue wet weight is recorded for each animal. In the example given in Figure 2 (Prokai et al., 2003), OVX Sprague–Dawley rats received 200 μg/kg body weight HEDD or E1 in corn oil vehicle, subcutaneously, 2 h before the onset of the 1-h middle cerebral artery occlusion. Control animals received the vehicle only. Rats were then euthanized 24h after reperfusion, the uteri were collected, and their weights were recorded.

**Figure 2.**
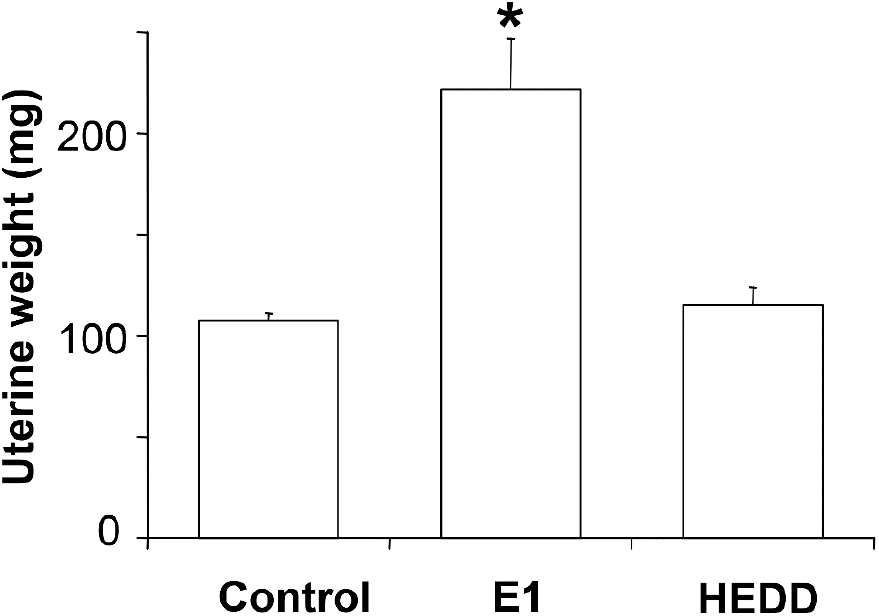
HEDD does not bind to ERα; therefore, it does not stimulate the rat uterus unlike its ER-binding parent estrogen, E1 (adopted from Prokai et al., 2003). Data are mean ± SEM, n=8-10. * p < 0.05 from control and HEDD, respectively.

### Docking Studies

The three-dimensional structure of ERα’s ligand-binding domain complexed to E2 was obtained from the Protein Data Bank (http://www.rcsb.org/pdb/, entry number: 1A52). After visualization and preparation for molecular docking with HEDD using the SeeSAR software suite (version 11.0.2; BioSolveIT GmbH, Sankt Augustin, Germany), AutoDock Vina and AutoDock Tools (Trott and Olson, 2010) were used to perform the modelling experiments.

## Acknowledgement

Work from our laboratory included in this Letter to the Editor was generated by the financial support of the National Institutes of Health, Bethesda, MD, USA (grant numbers AG031421, EY027005 NS044765, AG031535, AG031387, MH100700, HD078077 and CA215550), as well as by the Robert A. Welch Foundation (endowment BK-0031). The authors thank Daniel L. De La Cruz for reviewing the computational studies reported by Canário et al. (2022).

